# Complexity analysis of heartbeat-related signals in Brain MRI time series as a potential biomarker for ageing and cognitive performance

**DOI:** 10.1101/2020.05.27.117226

**Authors:** David López Pérez, Arun L. W. Bokde, Christian M. Kerskens

**Affiliations:** Institute of Psychology, Polish Academy of Sciences, Warsaw, Poland; Institute of Neuroscience, Trinity College, Dublin, Ireland; Trinity College Institute of Neuroscience and Cognitive Systems Group, Discipline of Psychiatry, School of Medicine, Trinity College Dublin, Ireland

## Abstract

Getting older affects both the structure of the brain and some cognitive capabilities. Until now, magnetic resonance imaging (MRI) approaches have been unable to give a coherent reflection of the cognitive declines. It shows the limitation of the contrast mechanisms used in most MRI investigations, which are indirect measures of brain activities depending on multiple physiological and cognitive variables. However, MRI signals may contain information of brain activity beyond these commonly used signals caused by the neurovascular response. Here, we apply a zero-spin echo (ZSE) weighted MRI sequence, which can detect heartbeat evoked signals (HES). Remarkably, these MRI signals have properties only known from electrophysiology. We investigated the complexity of the HES arising from this sequence in two age groups; young (18-29 years) and old (over 65 years). While comparing young and old participants, we show that the complexity of the HES decreases with age, where the stability and chaoticity of these HES are particularly sensitive to age. However, we also found individual differences which were independent of age. Complexity measures were related to scores from different cognitive batteries and showed that higher complexity may be related to better cognitive performance. These findings underpin the affinity of the HES to electrophysiological signals. The profound sensitivity of these changes in complexity shows the potential of HES for understanding brain dynamics that need to be tested in more extensive and diverse populations with clinical relevance for all neurovascular diseases.

## 1 Introduction

Normal ageing has cascading effects on many cognitive domains, and it inflicts the brain at multiple levels ranging from sub-to macro-cellular (e.g., [1, 2]). For instance, older adults have particular difficulties with episodic memory [3], working memory (e.g., [4]) or are slower processing different stimuli (e.g., [5]). At the same time, some aspects of cognition are maintained, such as semantic memory ([6]) or emotional regulation [7]. However, the cerebral mechanisms that underlie this better or lesser performance are still poorly understood [8].

Many studies based on magnetic resonance imaging (MRI) have addressed how changes in the ageing brain may occur. Among the various MRI methods, functional MRI (fMRI), with more than 10,000 published papers, is probably the most widely applied [9]. The blood oxygen level-dependent (BOLD) signal obtained from fMRI is an indirect index of neural activity reflecting the neurovascular response. It depends on multiple variables that can alter due to physiological, pathological or psychological factors. Besides, the BOLD signal mainly derives from venous blood, reflecting the integrated flow and metabolism changes during the transit through the brain. Hence, fMRI studies in ageing research may deliver inconclusive findings. In this respect, studies have shown no clear direction. The majority of studies have reported that responses have been similar in both groups [8], but in some cases, the magnitude of the BOLD response was reduced in older adults [11], while in others, it was increased [12]. Reductions are often interpreted as cognitive deficits [13], while the increase is often considered compensatory [14] or a reduction in the selectivity of responses [15]. Therefore, ageing processes are still not completely understood, and a complementary method, which could target other aspects of the brain dynamics, would be desirable.

The heartbeat, essential for the brain to function, is rarely considered directly responsible for cognitive changes. This comes as a surprise because heart functions also alter with age, which should, in turn, affect cerebral blood flow. It is well known that several heartbeat related effects influence conscious perception, where the cardiac cycle may impact the perception of visual or auditory stimuli (e.g., [16]). The existence of heartbeat-evoked potential (HEP) in general is strong evidence that the heartbeat influences neuronal functions [17]. However, neither the origin of HEPs nor their relation to ageing is clearly understood, partly due to the low signal intensity in EEG or MEG and partly due to our poor understanding of electrophysiology in general. A recent study by Kerskens and López Pérez [18] may shed some light on the heart-brain relationship. They discovered heartbeat evoked signals in fast MRI time series, which resembled some properties of HEPs; (a) they appeared in a similar time window at 300 ms after cardiac R-wave, and (b) they were only visible if the volunteers were awake. Hence, MRI may allow studying the origin of some electrophysiological phenomena which, in contrast to the BOLD signal, are directly related to brain activity. Combined with the advantage that MRI signals can be localised, those signals may open up new opportunities. However, the contrast mechanism behind those heartbeat evoked signals (HES) is still unknown. Electromagnetic fields of brain cells [20] as well as any known MRI contrast including inflow, BOLD, T2/T1 relaxation, magnetisation transfer or diffusion [18] could be excluded as potential mechanisms. Moreover, Kerskens and López Pérez showed that MRI signals can, despite common believe [23], behave non-classically, which is in accordance with earlier controversial ideas connecting brain functioning with quantum mechanics [25]. Recently, some studies have indicated that quantum coherence in living organisms may exist and be essential for their functioning (e.g., [26, 27]) as well as influence brain activity and affect cognition [28]. Therefore, even without interpretational consent, HES are interesting for further explorations.

So far, we know that HES show features of dipole-dipole (or spin-spin) interactions. Those interactions can be derived classically, referring to multiple spin echoes (MSE) or quantum mechanically. The quantum mechanical derivation in fluids is commonly referred to as intermolecular multiple quantum coherence (iMQC or just MQC). This naming convention is somewhat misleading because of the equivalence to the classical derivation [21], which indicates that the MRI signal contains no quantum features. Therefore, MQC has not traditionally been considered a powerful tuning element for enhancing or explaining functions in biology [24]. Also, due to its low signal-to-noise ratio, MSE/iMQC sequences have not been considered for fast time series either. However, the use of fast time series results in saturated signals, in quantum mechanical terms “mixed quantum states”, which are necessary to detect those heartbeat evoked signals [18]. For that reason, those signals deviate from the classical MSE/iMQC signals. A further investigation of the neuronal basis may help to shed light on the underlying contrast mechanisms. In this respect, it has been shown that the HES were robust and reliable to reproduce [18]. Nevertheless, HES showed high variability and complexity, suggesting that the interaction between the brain and the heart and its complexity would be high-dimensional [28, 29].

Recent studies have shown that high complexity is characteristic of healthy systems and they can degrade because of disease or ageing [30]. Thus, if this mechanism is vital for cerebral dynamics, the complexity of these fluctuations needs to be high and any variation in the dynamics with age should affect the complexity of the system. Here, we want to investigate, for the first time, the HES in two age groups (one between 18-29 years and the other +65 years) and study how the dynamical complexity of the signal varies in each of them. We use a broad range of dynamic systems measures to characterise these fluctuations as entirely as possible. First, we applied Recurrence Quantification Analysis (RQA; [31]), which is an increasingly popular method to analyse dynamic changes in behaviour in complex systems. This concept has been used to study physiological signals [32, 33], heart rate variability [30] or the dynamics of heart rhythm modulation [34]. The main benefits of RQA compared to standard analysis reside in its sensitivity to small changes in the system dynamics [30]. Secondly, we employed MultiFractal Detrended Fluctuation Analysis to extract the fractal properties of the signal (MFDFA; [35]). Multifractal Analysis is another efficient chaos theory method to study the fractal scaling properties and long-range correlations of noisy signals [29] (for a review see [36]). Differences if fractal measures as a consequence of ageing have been found in electroencephalography (EEG) [37] or due to increased heartrate variability (HRV) changes [38]. Finally, we relate these measures with different cognitive batteries and show that those quantum fluctuations may be essential for cerebral dynamics and cognitive functioning.

## 2 Methods

### 2.1 Participants

60 subjects (29 participants between 18 and 29 years old, and 31 participants over 65 years old) were scanned with the protocols approved by the St. James Hospital and the Adelaide and Meath Hospital, incorporating the National Children Hospital Research Ethics Committee. All participants were adults recruited for a larger study [39–41] and came from the greater Dublin area. All participants underwent the Cambridge Neuropsychological Test Automated Battery (CANTAB; [42]) which has been used to detect changes in neuropsychological performance and include tests of working memory, learning and executive function; visual, verbal and episodic memory; attention, information processing and reaction time; social and emotion recognition, decision making and response control. The CANTAB scores were normalised for age and IQ. Particularly, the following subtest were administered:

- The Paired Associate Learning Test is a measure of episodic memory where boxes are displayed on the screen, and each one has a distinct pattern. The boxes are opened in random order, revealing the pattern behind the box. In the test phase, patterns are individually displayed in the centre of the screen, and participants must press the box that shields the respective pattern.
- Pattern Recognition Memory is a test of visual pattern recognition memory in which the participant is presented with a series of visual patterns, one at a time, in the centre of the screen. In the recognition phase, the participant is required to choose between a pattern they have already seen and a novel pattern. In this phase, the test patterns are presented in the reverse order to the original order of presentation. This is then repeated, with new patterns. The second recognition phase can be given either immediately (immediate recall) or after a delay (delay recall).
- The Spatial Working Memory Test assesses spatial working memory in which boxes are presented on the computer screen and hidden behind one of the boxes is a yellow circle. Participants must find the box where the yellow circle is located. As the task progresses, the number of boxes on the screen increases. We analysed the spatial working memory strategies (i.e., the number of times participants begin a new search strategy from the same box).

Moreover, participants performed the trail making test (TNT; [43]) which is a neuropsychological test of visual attention and task switching. TNT test that can provide information about visual search speed, scanning, speed of processing, mental flexibility, as well as executive functioning [44].

### 2.2 Data Acquisition

Each participant was imaged in a 3.0 T Philips whole-body MRI scanner (Philips, The Netherlands) using a standard single-shot Gradient-Echo Echo Planar Imaging (GE EPI) sequence operating with a 8-channel array receiver coil in all cases. The parameters of the EPI time-series sequence were as follows: Radiofrequency flip angle = 30^°^, repetition time (TR) = 60 ms and the echo time (TE) = 18 ms with a voxel size was 3.5 × 3.5 × 3.5 mm, matrix size was 64 × 64, SENSE factor 2 (reduction factor of the phase encoding steps compensated by receiver coil design), bandwidth in readout direction was 2148 Hz (data sampling rate). In addition, two saturation slices of 5 mm in thickness were placed parallel to the imaging slice (15 mm above and 20 mm below; see Figure 1). These slabs were applied to introduce asymmetrical magnetic field gradients (labelled with “s” in Figure 2) of 6 ms duration and 21 mT/m gradient amplitude between two EPI scans which leads to condition that can generate zero quantum coherence (sZQC; gradients labelled with “r” and “c” in Figure 2 also contributed to the asymmetry). As a result, the sequences time series produced alternating sZQC. The signal amplitude depends on the slice orientation and changing the slice orientation varies the demagnetization field. The duration and strength of the gradients influence the correlation distance *d* (see Supplementary Figure 4 for a full description see [18,49]). The pulse sequence scheme indicating most of the parameters above is shown in Figure 2. The imaging slice was set coronal above the ventricle to avoid pulsation effects (see Figure 1). The average angulation of the imaging slice was 14.76 ± 5.65 degrees. The angulation was in accordance with the other sequences in the imaging protocol. It was 5 ± 5 degrees off the optimum magnetic field directions, which reduced the sZQC effect only slightly.The total duration of the scan was 1 minute.

**Fig. 1.**
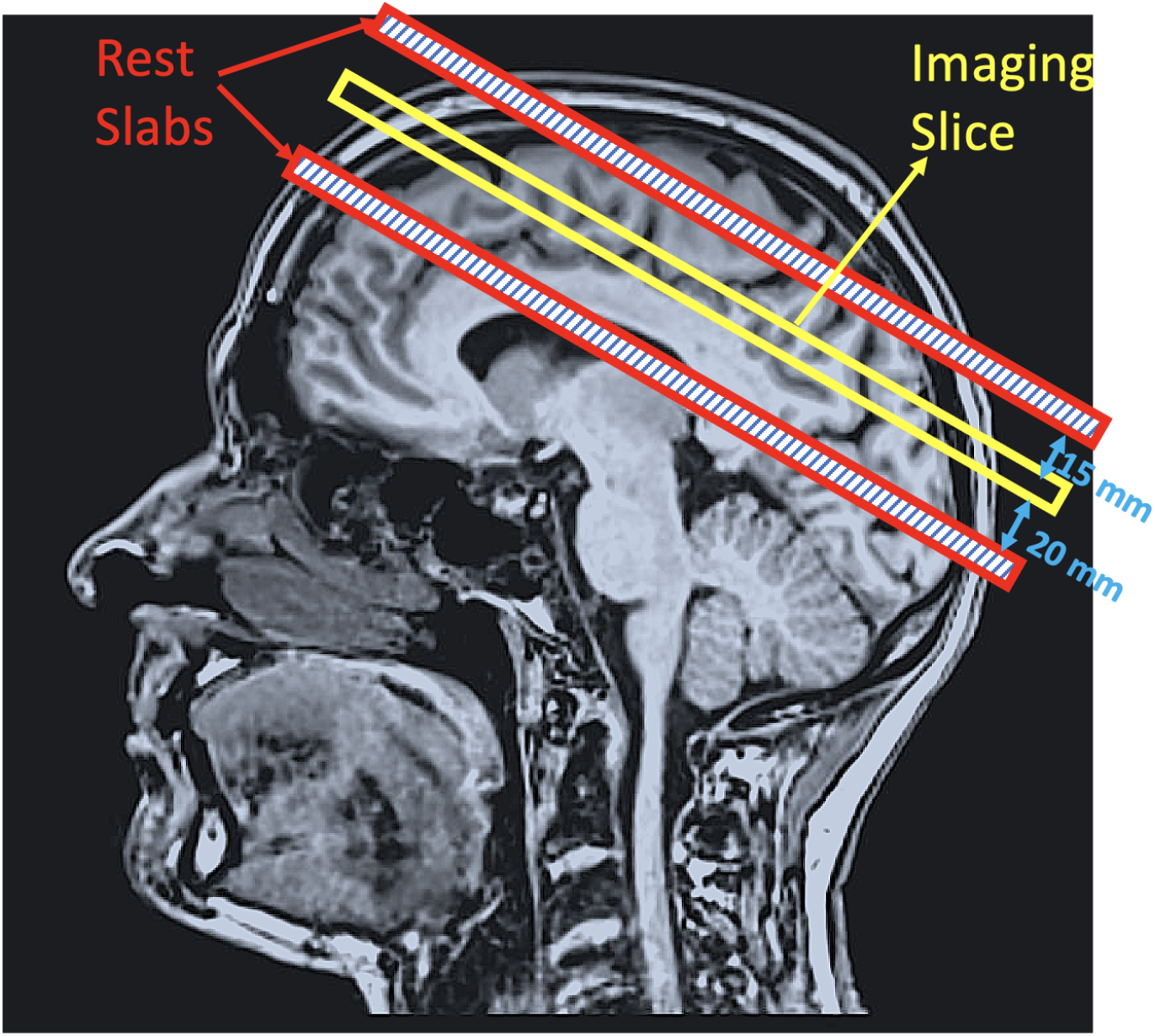
Example of the acquisition model which includes the imaging slice (central red line) and the REgional Saturation Technique (REST) slabs above and below the imaging slice both 5mm thick and separated 15mm and 20mm respectively.

**Fig. 2.**
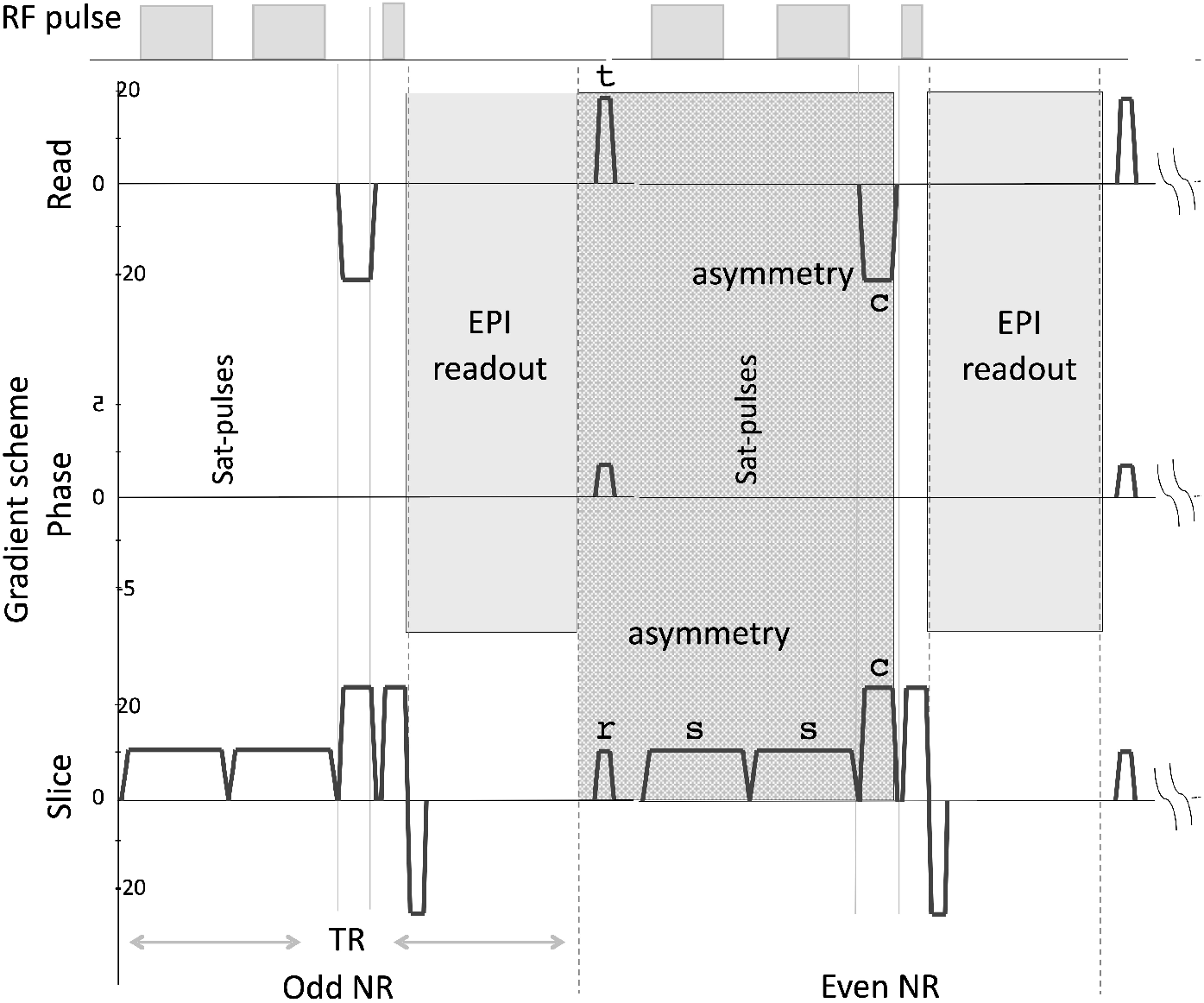
MRI acquisition sequence.

**Fig. 3.**
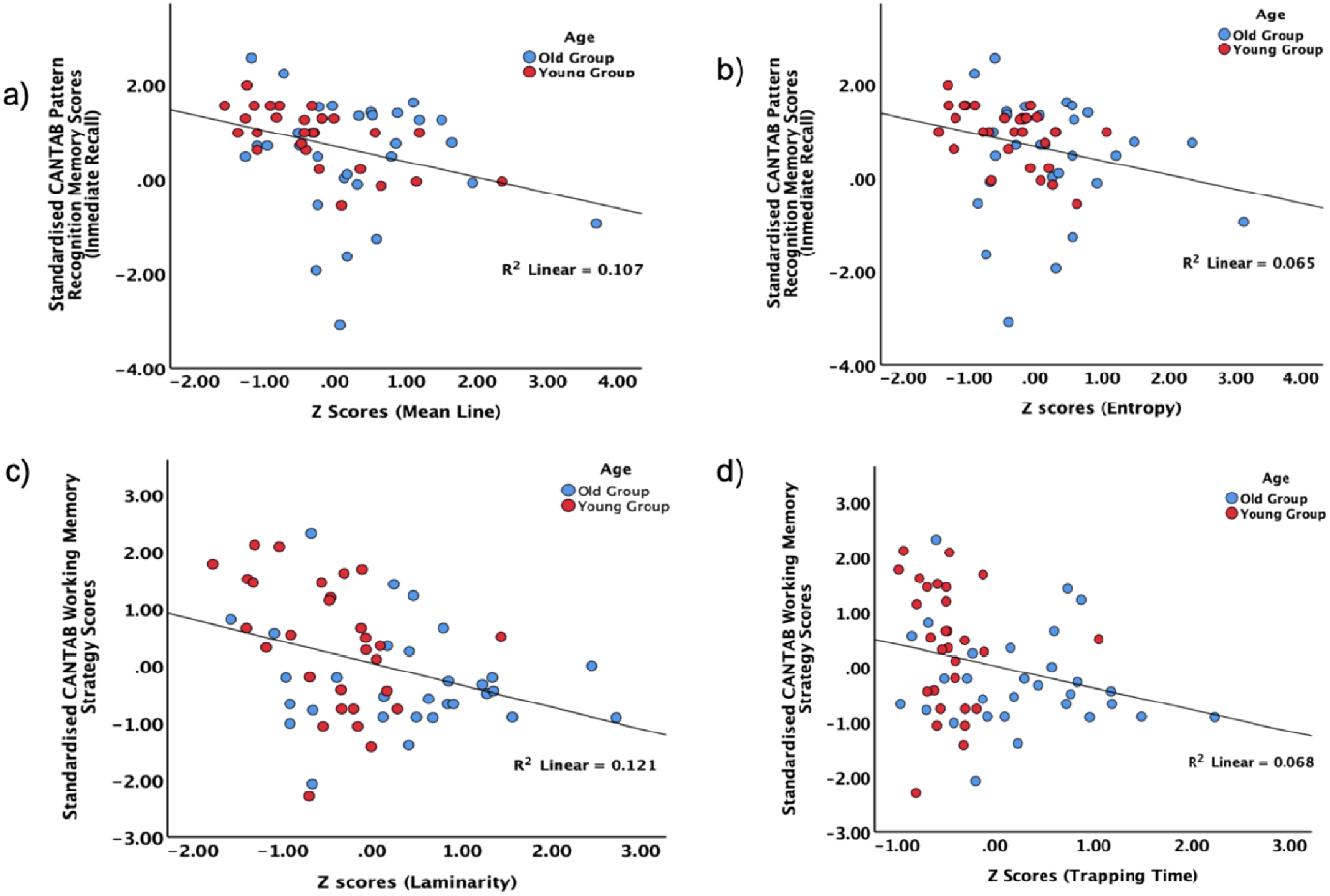
Examples of linear regressions between mean line (a) and entropy(b), and Standardised CANTAB Pattern Recognition Memory Scores (Inmediate Recall), and between Laminarity (c) and Trapping Time (d), and standardised CANTAB working memory strategy scores.

Anatomical MRI images in all studies included a high-resolution sagittal, T1-weighted Magnetization-Prepared RApid Gradient-Echo (MP-RAGE) (TR = 2.1 s, TE = 3.93 ms, flip angle = 7^°^). The sequence was acquired after the resting-state fMRI part of the session. The radiographer always contacted the participants before the acquisition to make sure that they were awake. This step is important given that HES are sensitive to changes in wakefulness of the participant [18].

### 2.3 Signal Pre-processing

All calculations were developed in a Dell Optiplex 790 with 12 Gb RAM using Matlab 2017a (The MathWorks Inc., Natick, MA, 2017). The MRI time series was calculated as the average signal across all the voxels in the imaging slice. Since motion correction could not be applied due to the single slice nature of the experiment, average time-series were visually inspected in search for irregularities which were manually removed from the analysis leaving the rest of the time-series unaltered. In addition, the data was not smoothed to avoid removing high frequencies which may lead to the loss of information [45]. Manual segmentation was used to create a mask to remove cerebrospinal fluid (CSF) contributions which were later eroded to avoid partial volume effects at the edges. The first 100 scans were removed to avoid signal saturation effects

### 2.4 Recurrence Quantification Analysis

We used Recurrence Quantification Analysis (RQA) to analyse the dynamical temporal characteristics of the MRI signals. RQA quantifies the repeated occurrences of a given state of a system (i.e., recurrences) by analysing the different structures present in a recurrence plot (RP), which is a graphical representation of the recurrences in the dynamical system [31]. For a MRI time series *x* and radius parameter *ϵ* we define recurrence as:

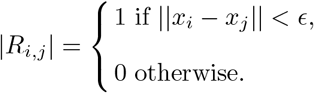

where *i* and *j* represent each time point of the time series *x* with length N. The result is a NxN RP of 1s and 0s which contains all the recurrent events in *x*. A series of variables describing the dynamics can be obtained from this plot [31]. In our analysis, we considered the following RQA measures [46]:

- Determinism (*Det*): it represents a measure that quantifies repeating patterns in a system, and it is a measure of its predictability. *Det* is defined as follows:

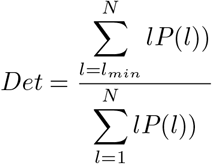

where *l* is the length of a diagonal line present in the RP, *P* (*l*) is the distribution frequency of diagonal line lengths, *N* is the maximum diagonal line length, and *l*_*min*_ is the minimum diagonal line length. Regular, periodic signals, such as sine waves, have higher *Det* values, while uncorrelated time-series cause low *Det*.
- Mean Line (*MeanL*): it is the average length of repeating patterns in the system and it is calculated as:

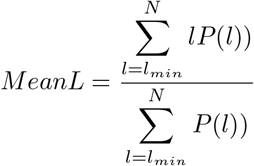

It represents the mean prediction time of the signal, a measure of chaos or divergence from an initial point.
- Entropy (*Ent*): it is the Shannon entropy of the distribution of the repeating patterns of the system and it is calculated as:

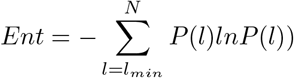

If a signal has high entropy it exhibits diversity in short- and long-duration periodicities.
- Laminarity (*Lam*): it determines the frequency of transitions from one state to another, without describing the length of these transition phases. *Lam* is computed as:

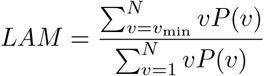

where *v* is the length of a vertical line present in the RP, *P* (*v*) is the distribution frequency of vertical line lengths, *N* is the maximum vertical line length and *v*_*min*_ is the minimum vertical line length. It indexes the general level of persistence in some particular state of one of the time-series [47].
- Trapping Time (TT): it represents the average time the system remains on a given state and it is a measure of the stability of the system:

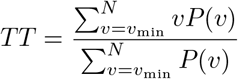

It was calculated here using the tt function from the Cross-Recurrence Plot (CRP) Toolbox for [48].
- Maximum Line (*MaxL*): it is the largest Lyapunov exponent of a chaotic signal, which gives the longest time spent in a single state by the system [50]. It can be calculated as:

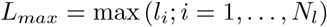

Three critical parameters need to be set to calculate the recurrence plots. First, the smallest sufficient embedding dimension was determined using the *fnn* function [51] within the CRP Toolbox (Marwan, N.: Cross Recurrence Plot Toolbox; [52], for MATLAB, Ver. 5.22 (R31.2), http://tocsy.pik-potsdam.de/CRPtoolbox/). This function estimates the minimum embedding dimension where the false nearest neighbours vanish. We applied the *fnn* to all time-series and obtained an average value of 15, which agrees with the typical values recommended for biological signals [32]. The second parameter is the delay which we calculated using the *mi* function from the CRP Toolbox [52, 53]. This function finds the non-linear interrelations in the data and determines which delay fulfils the criterion of independence. In the same way as the embedding dimension, we applied the mi function to all time-series and we obtained an average value of 3. Finally, several criteria have been suggested for the choice of the recurrence threshold [54]. Here, we adapted the radius for each time-series using the embedding dimension and delay computed together with a recurrence rate sufficiently low (i.e., RR = 3%) [48]. Additional parameters in the RQA calculations were Euclidean normalisation for each time-series and minimum line length equal to 2.

### 2.5 Multifractal Detrended Fluctuation Analysis

In biological systems, the coupling between different systems often exhibits [29] different spatial and temporal scales and hence its complexity is also multi-scale and hierarchical [29]. Moreover, biomedical signals often display temporal and spatial scale-invariant properties that indicate a multifractal structure. These structures can be defined by a multifractal spectrum of power-law exponents whose properties might be quantified with a few variables [35]. For instance, changes in the multifractal spectrum have been related to the variation with age or disease of the scale-invariant structures of some biomedical signals (e.g., [38]). Thus, to analyse the scale-invariant properties of the MRI segments and their changes with age, we used Multifractal Detrended Fluctuation Analysis (MFDFA). We first calculated the multifractal spectrum, D, of each time series using the MFDFA Matlab Toolbox [35]. The multifractal spectrum identifies the deviations in fractal structure within time periods with large and small fluctuations [35]. Each spectrum was computed using a window length with a minimum value of 2 and a maximum value of half the length of the time series. The q-order statistical moments were chosen between -11 and 11 and divided into 21 steps (see further description in [35]).

Two variables were calculated from each fractal spectrum, i.e., the width of the spectrum *W* and the position of the spectrum maxima *H*. The width *W* is calculated by subtracting the lower part of the spectrum, *h*, from the upper part of the spectrum, *h* [35, 38, 55]:

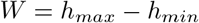

where a small width indicates that the time-series has fewer singularities and tends to be more monofractal. Finally, the *H* variable represents the value *h* in which the singularity spectra has its maximum [38]:

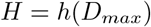

The position of *h* moves to higher values when the stronger singularities are present. Highly deterministic signals can often be explained by a lower number of fractal dimensions and are characterised by smaller *W* and *H* due to a decrease in the number of singularities.

### 2.6 Statistical Analysis

Before any statistical analysis, all variables were converted to *z* -scores. Those participants having *z* -scores larger than 3 standard deviations in three non-linear variables or more were rejected from the analysis. In total only 1 participant in the old group was removed. Independent *t* -tests were applied to compare the results. The participants grouped by age (young or old) were the independent variable and their RQA and Fractal variables were the dependent variables. Given the number of variables calculated, methods like logistic regression might be a better choice to find differences between both age groups. However, RQA and fractal measures tend to be highly correlated within each type of nonlinear analysis. This complicates the inclusion of any of the variables in any regression model since the model does not survive a second step and just one variable is enough to explain the variance between groups. Although highly correlated, these variables measure different things, and we thus chose independent t-tests to search for differences between both age groups. However, we additionally created a logistic regression model with all the RQA and fractal measures as predictors and age group as a dependent variable to test the accuracy of these measures to differentiate between age groups(see Supplementary Materials for the full results of the logistic regression analysis). Inspection of Q-Q Plots was carried out to all the measures to check if the data were normally distributed. Additionally, Levene’s test for equality of variances was applied and, in those cases, where this assumption was violated, a *t* statistic not assuming homogeneity of variance were computed on these measures. Finally, Spearman’s correlations between the RQA variables and the cognitive batteries were performed.

## 3 Results

### 3.1 Non-linear dynamics of the Heart-Evoked Signals

First, we tested how the non-linear dynamics of the HES varied with age across all participants. Significant correlations with age were found for *Lam* (*r* (59) = .228, *p* < .001) or *TT* (r(59) = .29, *p* < .001). Additional correlations were *Det* (*r* (59) = .03, *p* = .18), *MeanL* (*r* (59) = .08, *p* < .02), *MaxL* (r(59) = .11, *p* < .029) and Ent (r(59) = .12, *p* < .03) (see also Supplementary Figures 1 and 2). At a group level all the RQA variables but the DET were statistically significantly higher in the old group in comparison to the young one (see Table 1 for group averages): MeanLine (*t* (57) = 2.23, *p* = .02; d = .58), MaxLine (*t* (57) = 2.81, *p* = .007; d = .73), Ent (*t* (57) = 2.62, *p* =.01; d = .68), Lam (*t* (57) = 3.68, *p* = .001; d = 0.96) and TT (*t* (57) = 4.57, *p* < .001; d = 1.19) and Det (*t* (57) = 1.23, *p* = .22; d = .32).

**Table 1.**
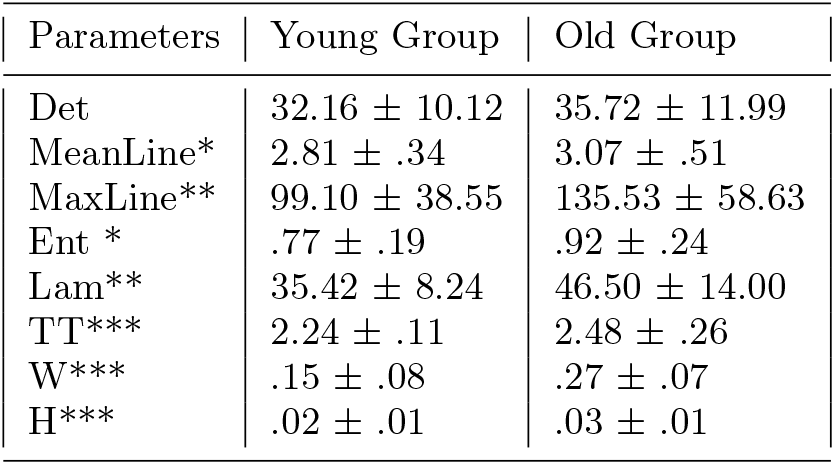
*Group mean averages of the RQA and MDFDA variables* extracted from the ZQC weighted time-series for the young and old groups (*p* < .05(*),*p* < .01(**),*p* < .001(***))

Likewise, Significant correlations with age were found for *W* (*r* (59) = .343, *p* < .001), *H* (*r* (59) = .183 (see also Supplementary Figures 1). The fractal properties of the HES in the old group were statistically higher in W (*t* (57) = 5.44, *p* <.001; d = 1.41) and H (*t* (57) = 3.53, *p* < = .001; d = .92) in comparison to the young group, suggesting a more chaotic behaviour in the old population.

Finally, we run a logistic regression model with all the RQA and Fractal as predictors and age group as a dependent variable. The model was significant (*χ*^2^(8) = 38.57, *p* < .001) and both groups were classified with an overall accuracy of 86.4%,(see Supplementary Analysis for full logistic regressions results). This highlights the sensitivity of the HES to changes in age.

### 3.2 Are group differences coming from movement or cognitive differences?

Since the HES are sensitive to movement, we explored the relationship between the non-linear variables and motion quality control variables from the rs-fMRI as a proxy for potential average movement of the participant Although the information was not available for the young group, there were no significant correlations between these measures (see Table 2) or in other words, the non-linear dynamics in the old cohort were not worsened by motion.

**Table 2.** *Spearman Correlations between quality control measures from the rs-fMRI session and the non-linear variables* from the ZQC signals. In these correlations, n was equal to 27 since movement information was not available for the young group, one participant did not have movement information and another did not pass quality control (see Section 2.5)

| Variables | QC max movement | QC mean movement |
| --- | --- | --- |
| Det | -.36 (.07) | .13 (.50) |
| MeanLine | -.10 (.60) | .00 (.97) |
| MaxLine | -.30 (.13) | -.01 (.95) |
| Ent | -.26 (.19) | .08 (.69) |
| Lam | -.27 (.17) | .17 (.38) |
| TT | -.13(.52) | .22 (.27) |
| W | .06 (.77) | .02 (.92) |
| H | -.14 (.47) | -.06 (.77) |

Finally, we explored any possible relation to cognitive measures and tests performed during the study. CANTAB scores showed consistent negative correlations and trends (see Table 3, Figure 2) between visual memory scores (pattern recognition memory and working memory) and the RQA variables while no correlations arose with the TNT scores. Consistently, young participants that showed higher complexity (i.e., smaller non-linear variables) also had better cognitive scores. Interestingly, age only correlated in only one of tests(Pattern Recognition Mem. (delayed recall): *r* (59) = -.25, *p* < .05) while no relations arose in the other tests (Pat. Recognition Mem. (immediate recall): *r* (59) = -.13, *p* = .29, Paired Associates Learning: *r* (59) = -.00, *p* = .94 ; Spat. working Mem. (Strategy): *r* (59) = -.14, *p* = .27; Trial A: *r* (59) = .20, *p* = .13; Trial B: *r* (59) = .18, *p* = .18). Altogether, there were significant changes in complexity with age and those changes were related to some cognitive scores, which in turn did not show any age effect. This suggests that the complexity changes of the HES may be sensitive to some aspects of cognition.

**Table 3.** Spearman correlations between the non-linear variables of the ZQC signals and the CANTAB and TNT scores. In these correlations, n varies between 59 (CANTAB scores) and 55 (TNT scores) since one participant did not pass quality control (see section 2.5) and the measures were not available to all of them. p-values are in parenthensis (trend (+), p <.05(*) and p <.01(**)).

| Variables | Pat. Recognition Mem. (immediate recall) | Paired Associates Learning | Spat. working Mem. (Strategy) | Pattern Recognition Mem. (delayed recall) | Trial A | Trial B |
| --- | --- | --- | --- | --- | --- | --- |
| Det | -.31 (.01)* | .06 (.62) | -.19 (.13) | -.02 (.87) | .05 (.69) | -.02 (.87) |
| MeanLine | -.33 (.009)** | -.04 (.71) | -.17 (.17) | -.13 (.31) | .00 (.98) | -.04 (.74) |
| MaxLine | -.34 (.008)** | -.18 (.16) | -.26 (.04)* | -.19 (.12) | -.00 (.96) | .02 (.86) |
| Ent | -.33 (.009)** | .06 (.65) | -.21 (.10) | -.12 (.33) | .09 (.49) | -.04 (.79) |
| Lam | -.29 (.02)* | .03 (.78) | -.34 (.008)** | -.09 (.48) | -.04 (.72) | -.05 (.68) |
| TT | -.25 (.05)+ | -.08 (.51) | -.29 (.02) | .18 (.15) |  |  |
| W | -.18 (.14) | -.04 (.71) | -.13 (.29) | -.17 (.17) |  |  |
| H | -.24 (.06) | .04 (.75) | -.22 (.10) | -.09 (.48) |  |  |

## 4 Discussion

In this paper, we have analysed the dynamical complexity of the heartbeat evoked signal (HES) in brain tissue of two age-groups. We found quantitative differences between both age groups, which were related to variations in complexity and chaos of the measured signals. Those variables, which showed the highest correlation (*TT,W, H*), had previously been reported as the strongest indicators of wakefulness/conscious awareness [18]. Additionally, we found that variables, which were less correlated with age, showed correlations with some CANTAB scales. A higher complexity of the signals was related to better cognitive performance, which in contrast, were not correlated to age. Interestingly, the CANTAB scales were related to short-term memory, which underpins the hypothesis further that the MR signals are of similar origin as HEPs.

In an earlier study, Kerskens and López Pérez reported that body movement introduced by hyperventilation can also reduce the signal amplitude. At the same time, hypoventilation which increases cerebral pulsation and blood flow has no qualitative effect on the signal. From this findings, we concluded that below a certain threshold HES may not be effected by motion [18]. In accordance with qualitative assessments of the time course (see Supplementary Figure 3 for examples), where we saw a signal amplitude decline in the 65+ group, we investigated if movement or pulsatile motion due to brain atrophy [56] could have influenced our results. Although the information was not available for the young group, motion quality control variables from an fMRI study within the same session did not correlate with any non-linear variables (see Table 2). Thus, we can conclude that motion was not significantly relevant. Regardless of this, future studies using the sequence should try to minimise the effect of movement during the data acquisition (e.g., adding extra cushions to hold the head), which might help improve the intensity of the signals [18].

Next, we studied how the HES declines with changes in age. To check that, we first quantified the apparent differences between both groups using non-linear timeseries analyses to determine changes in the dynamics of the MRI signals. First, we applied Recurrence Quantification Analysis (RQA), which was proven to be sensitive to small changes in the system dynamics and a powerful discriminatory tool to detect significant differences between both age groups. All the RQA measures (see Table 1) were lower in the young group, suggesting differences in the complexity of the underlying signal dynamics in both populations. Second, we applied fractal analysis to study the fractal scaling properties and long-range correlations of the signals. We showed an increase in the number of singularities with age, which is characterised by an increase in the width and position of the spectral maxima [35]. These differences were supported by the RQA entropy which denotes the Shannon entropy of the histogram of the lengths of diagonal segments and thus indicates the complexity of the deterministic structure of the system [30]. What is more, logistic regression analysis showed the predictability power those complexity measures have, with 86% of all participants classified accurately. Altogether, those results are in line with recent studies indicating that higher complexity in a system is a feature of healthy dynamics [30] or a higher degree of functional specialisation and integration in brain dynamics [57] and that this complexity declines with disease and age [58]. However, although there were group differences in complexity and most of the variables correlated with age, only half of them survived multiple significance corrections (i.e., *LAM, TT, W* and *H*). Those measures in particular, are related to the stability and chaoticity of a system, suggesting that more chaotic and stable behaviour (i.e., less complex) are characteristic of a decline with age. Thus, these complex measures might be used to spot differences in the underlying dynamics of the HES. Further testing acquiring physiological variables (e.g., blood pressure, heart rate and breathing) is needed to understand the origin of these HES.

Those variables which were not strongly correlated with age showed significant negative correlations with some CANTAB scores (see Table 3). Similarly to other studies showing that lower values are related to healthy dynamics (e.g., [35,59]), we found that lower scores (i.e., higher complexity) were related to better cognitive scores. This underpins the hypothesis that the HES are similar to HEPs and related to some aspects of cognition. Particularly, the relations with pattern recognition memory and working memory subscales suggest a link between the HES and short-term memory abilities. A potential explanation for why the HES was correlated to pattern recognition memory and spatial working memory is that the acquisition slice was roughly located in parietal and posterior cingulate regions. These are areas associated with these cognitive domains [60, 61]. Paired associate learning, however, is a hippocampus-based task [62]) and therefore, one would not expect to find a correlation with the measured signal. Besides, fMRI studies have shown that healthy old adults present higher activity levels in some brain regions during the performance of cognitive tasks, and these changes coexist with disrupted connectivity (for a review, see [63]). However, to the best of our knowledge, there are no fMRI-based signals that can predict these CANTAB scores consistently. This is especially surprising since the HES represents the average over the imaging slice and is a very rough and functional measurement. More importantly, the pattern recognition memory and working memory subscales which were strongly correlated with the non-linear variables, did not correlate with age. This finding emphasises the sensitivity of the HES towards cognitive changes. This is in line with the relation of the sZQC signal to conscious awareness, which Kerskens and Lopez Perez [18] reported earlier. The highly synchronisation over the imaging slice indicates that the signal origin could be a global physiological effect which may be essential for understanding the underlying brain computations.

However, despite the promising results, several limitations arise in this study. First, the acquisition protocol to obtain HES required fast repetition times, limiting the number of imaging slices to just one. The use of one imaging slice complicates the study of particular areas. It could induce variability in the results across all the participants, even when the position of the imaging slice is carefully planned. Consequently, additional slices should be acquired to study a larger region. This would allow studying specific brain areas and quantifying movement. Some approaches could be used to overcome this limitation. For example, next-generation MRI systems can acquire three or more imaging slabs using Multi-band excitation [19] with the same time resolution. A second improvement can be achieved by increasing the number of channels in the receiver coil, allowing the acquisition of data with shorter repetition times and a better signal-to-noise ratio. Future research should focus on expanding the sequence protocol to cover larger brain areas that would allow the use of the sequence in a wide range of studies. Secondly, with the information available, we could not differentiate if the group effects originated from an age-related reduction of wakefulness or an age-related loss of conscious awareness. Even though the radiographer was checking that all volunteers were awake before the data acquisition, under these conditions (i.e., testing in a supine position inside a dark room with no specific instructions but to remain still) participants in the older group are more likely to feel sleepy, thus potentially affecting their wakefulness [18]. In future studies, this could be controlled with eye-tracking equipment or further stimulation. However, it also provides a new tool to observe the volunteers’ wakefulness, which could improve MRI studies in the future. Further, it allows studying if the effect is reversible by wakening up the participants or if it is a real loss in awareness or other age-related declines. Thirdly, there was no access to any heart rate or physiological cardiac measures at the time of the experiment, which could be helpful given the importance of these cardiovascular fluctuations. Finally, the group sizes in this study were small, and the results need to be considered preliminary. Further research is needed to confirm these findings.

## 5 Conclusion

We have provided evidence that HES exist in the brain and their complexity decreased with age. Consistent with the idea that higher complexity is related to healthier dynamics, we showed quantitatively that the decline of these fluctuations is related to a decrease in the complexity of the signal time series with age. Additionally, some aspects of the higher complexity were related to better cognitive performance, which, like the CANTAB results, were not age-related. Hence, HES can differentiate between brain functions that decline with age and those that do not. Altogether, the complex properties of the HES provide a potential mechanism for understanding brain dynamics that need to be tested in more extensive and diverse populations with clinical relevance for all neurovascular diseases.

## 6 Conflict of Interest

The authors have declared no conflict of interest

## 7 Data availability

Data and scripts used in the analyses are available upon request.

## 8 Acknowledgements

We would like to thank Elizabeth G. Kehoe and Dervla Farrell for assistance in acquiring the data, the Trinity College’s IT Research, Sojo Joseph for carrying out the imaging protocols for all participants and Edyta Stanaszek for reading earlier versions of the manuscript. This work was supported by Science Foundation Ireland (SFI-11/RFP.1/NES/3051), the Science Foundation Ireland Stokes Programme (07/SK/B1214a), from the European Regional Development Fund via the Interregional 4A Ireland Wales Programme 2007–2013 and Trinity College Institute of Neuroscience. This project has received additional funding from the Institute of Psychology, Polish Academy of Sciences.

## Notes

### Competing Interest Statement

The authors have declared no competing interest.

### Summary of Updates

This version is the revised manuscript after reviewer comments.

