## supplementary files for "Complexity analysis of heartbeat-related signals in Brain MRI time series as a potential biomarker for ageing and cognitive performance"

### Supplementary Figures

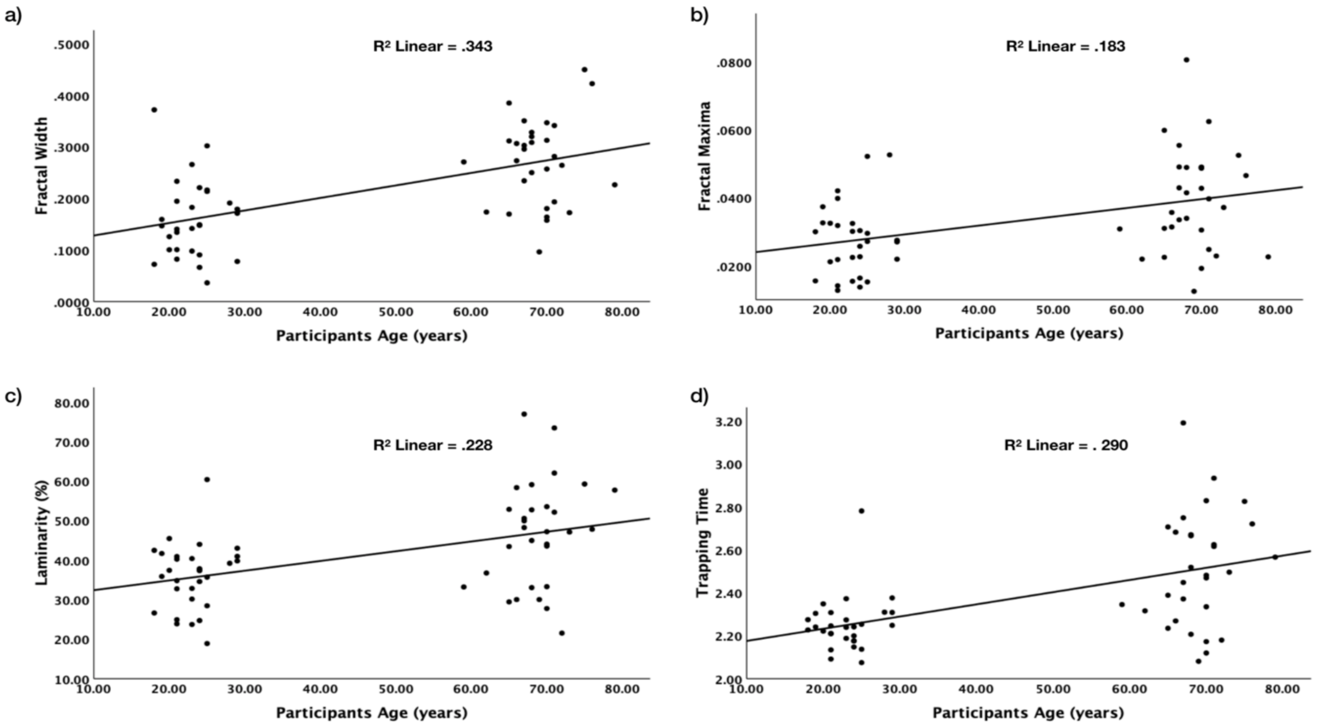


***Supplementary Figure 1.*** *Linear regressions between (a) fractal width, (b) fractal maxima, (c) laminarity and (d) trapping time, and participants age.*

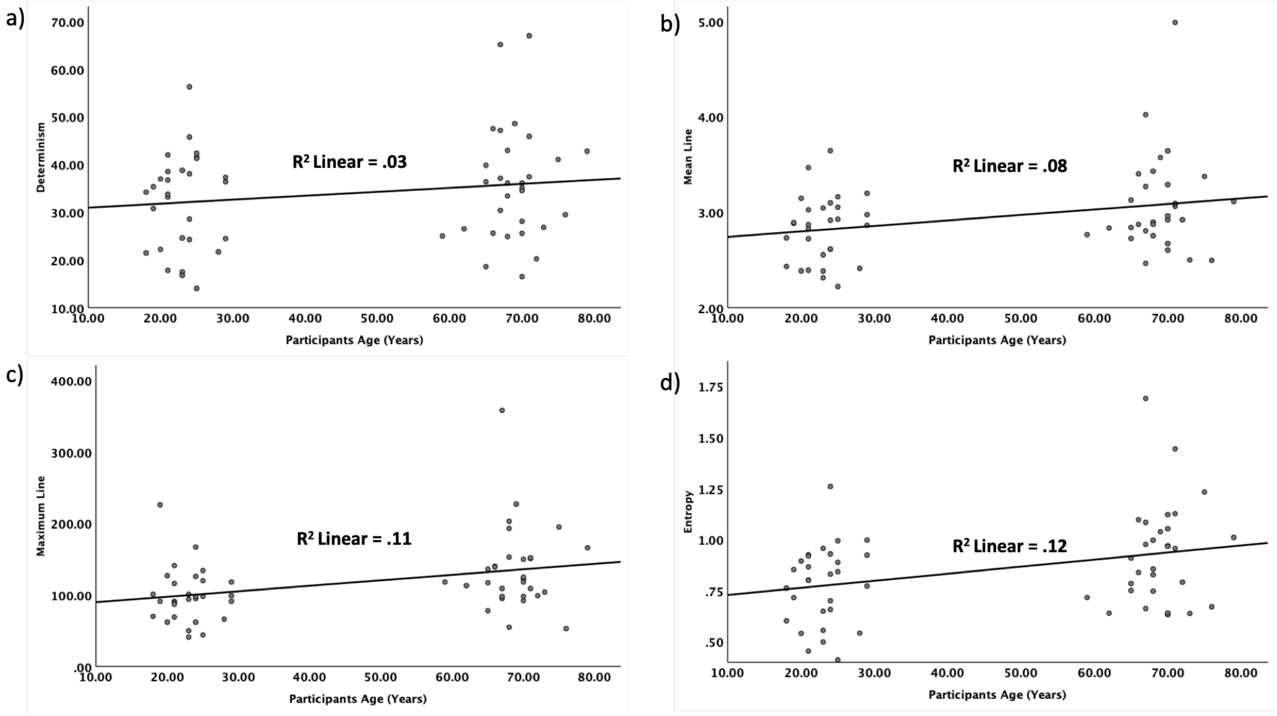


***Supplementary Figure 2.*** Linear regressions between (a) determinism, (b) mean line, (c) maximum line and (d) entropy, and participants age.

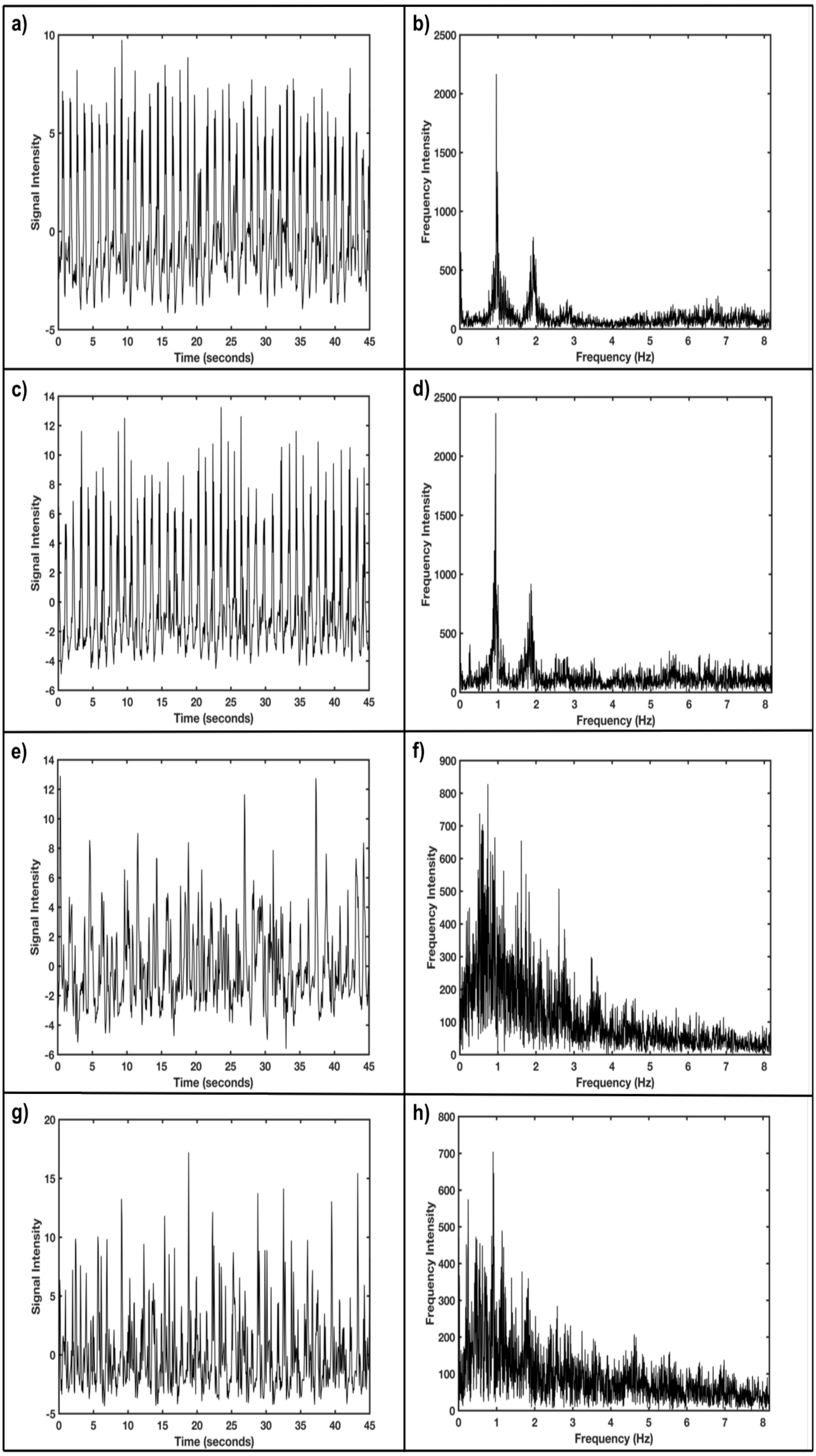


***Supplementary Figure 3.*** *Example of the time-series as well as its frequency spectra for two young (a-c and b-d) and two healthy old adults (e-g and f-h). These examples are representing the typical results in these groups.*

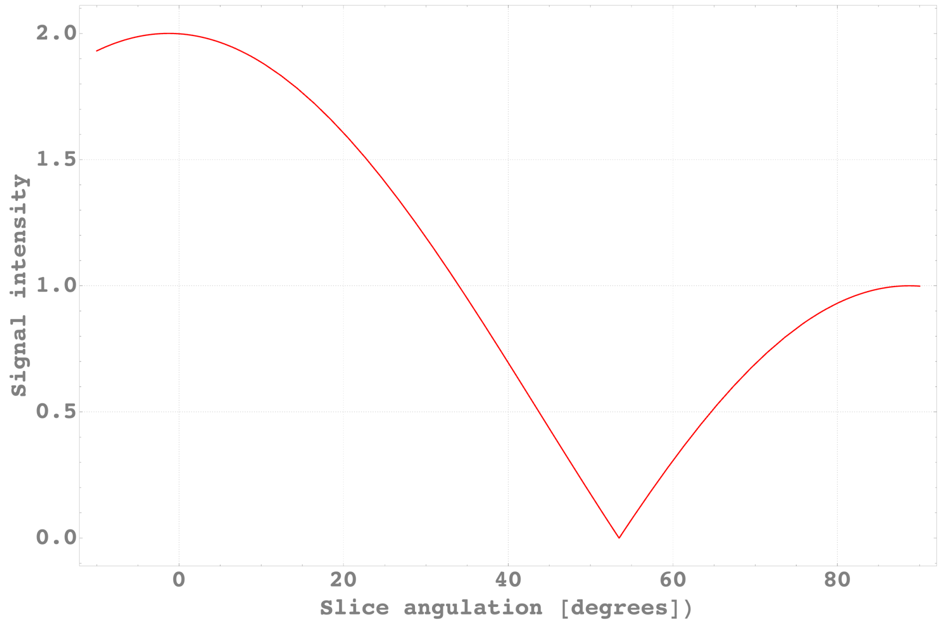


***Supplementary Figure 4.*** *Signal amplitude depending on the slice orientation. A minimum value is observed at the magic angle (aprox. 55 degrees).*

**Supplementary Analysis**

***Multiple Logistic Regression for the RQA parameters***

The logistic regression model with all the RQA variables in only survive the first step with TT being enough to classify both age groups.

**Variables in the Equation**

| | | B | S.E. | Wald | df | Sig. | Exp(B) |
|---|---|---|---|---|---|---|---|
| Step 1[a] | Zscore(TT) | 2.405 | .708 | 11.546 | 1 | <.001 | 11.075 |
| | Constant | .480 | .372 | 1.666 | 1 | .197 | 1.616 |

a. Variable(s) entered on step 1: Zscore(TT).

With the variables not added into the model because no model improvement was found when adding them.

**Variables not in the Equation**

| | | | Score | df | Sig. |
|---|---|---|---|---|---|
| Step 1 | Variables | Zscore(Det) | .144 | 1 | .704 |
| | | Zscore(MaxLine) | 2.401 | 1 | .121 |
| | | Zscore(MeanLine) | 3.141 | 1 | .076 |
| | | Zscore(Ent) | 2.724 | 1 | .099 |
| | | Zscore(Lam) | .299 | 1 | .584 |
| | Overall Statistics | | 14.082 | 5 | .015 |

The final classification table shows that both age groups can be classified with an accuracy of 78%

**Classification Table**[a]

| | | | Predicted | | |
|---|---|---|---|---|---|
| | | | Age | | Percentage Correct |
| Observed | | | .00 | 1.00 | |
| Step 1 | Age | .00 | 25 | 4 | 86.2 |
| | | 1.00 | 9 | 21 | 70.0 |
| | Overall Percentage | | | | 78.0 |

a. The cut value is .500

***Multiple Logistic Regression for the Fractals***

The logistic regression model with all the Fractals variables in only survive the first step with W being enough to classify both age groups.

**Variables in the Equation**

| | | B | S.E. | Wald | df | Sig. | Exp(B) |
|---|---|---|---|---|---|---|---|
| Step 1[a] | Zscore(fractalWidth) | 1.768 | .462 | 14.653 | 1 | <.001 | 5.859 |
| | Constant | .188 | .330 | .325 | 1 | .569 | 1.207 |

a. Variable(s) entered on step 1: Zscore(fractalWidth).

With the fractal maxima not added into the model because no model improvement was found when adding them.

### Variables not in the Equation

| | | | Score | df | Sig. |
|---|---|---|---|---|---|
| Step 1 | Variables | Zscore(fractalMax) | .465 | 1 | .496 |
| | Overall Statistics | | .465 | 1 | .496 |

The final classification table shows that both age groups can be classified with an accuracy of 74.5%

### Classification Table[a]

| Observed | | | Predicted | | |
|---|---|---|---|---|---|
| | | | Age | | Percentage Correct |
| | | | .00 | 1.00 | |
| Step 1 | Age | .00 | 22 | 7 | 75.9 |
| | | 1.00 | 8 | 22 | 73.3 |
| | Overall Percentage | | | | 74.6 |

a. The cut value is .500

***Multiple Logistic Regression for all the parameters included in the model***

Including all the variables in the model we get a classification accuracy of 86.4%, where the younger group

#### Variables in the Equation

| | | B | S.E. | Wald | df | Sig. | Exp(B) |
|---|---|---|---|---|---|---|---|
| Step 1[a] | Zscore(fractalWidth) | 1.707 | .709 | 5.799 | 1 | .016 | 5.510 |
| | Zscore(fractalMax) | .040 | .931 | .002 | 1 | .966 | 1.040 |
| | Zscore(Det) | -2.924 | 1.320 | 4.907 | 1 | .027 | .054 |
| | Zscore(MaxLine) | 1.427 | 1.930 | .546 | 1 | .460 | 4.164 |
| | Zscore(MeanLine) | .526 | .694 | .573 | 1 | .449 | 1.691 |
| | Zscore(Ent) | 2.699 | 2.352 | 1.317 | 1 | .251 | 14.869 |
| | Zscore(Lam) | -.388 | 1.515 | .066 | 1 | .798 | .678 |
| | Zscore(TT) | 2.014 | 2.108 | .912 | 1 | .339 | 7.491 |
| | Constant | .887 | .653 | 1.847 | 1 | .174 | 2.429 |

a. Variable(s) entered on step 1: Zscore(fractalWidth), Zscore(fractalMax), Zscore(Det), Zscore (MaxLine), Zscore(MeanLine), Zscore(Ent), Zscore(Lam), Zscore(TT).

#### Classification Table[a]

| Observed | | | Predicted Age .00 | Predicted Age 1.00 | Percentage Correct |
|---|---|---|---|---|---|
| Step 1 | Age | .00 | 26 | 3 | 89.7 |
| | | 1.00 | 5 | 25 | 83.3 |
| | Overall Percentage | | | | 86.4 |

a. The cut value is .500
